# Ancestry of the two subgenomes of maize

**DOI:** 10.1101/352351

**Authors:** Michael R. McKain, Matt C. Estep, Rémy Pasquet, Daniel J. Layton, Dilys M. Vela Díaz, Jinshun Zhong, John G. Hodge, Simon T. Malcomber, Gilson Chipabika, Beatrice Pallangyo, Elizabeth A. Kellogg

## Abstract

Maize (*Zea mays* ssp. *mays*) is not only one of the world’s most important crops, but it also is a powerful tool for studies of genetics, genomics, and cytology. The genome of maize shows the unmistakable signature of an ancient hybridization event followed by whole genome duplication (allopolyploidy), but the parents of this event have been a mystery for over a century, since studies of maize cytogenetics began. Here we show that the whole genome duplication event preceded the divergence of the entire genus *Zea* and its sister genus *Tripsacum*. One genome was donated, in whole or in part, by a plant related to the modern African genera *Urelytrum* and *Vossia*, although genomic rearrangement has been extensive. The other genome donor is less well-supported, but may have been related to the modern *Rottboellia*-*Hemarthria* clade, which is also African. Thus *Zea* and *Tripsacum* together represent a New World radiation derived from African ancestors.

## Significance statement

Maize is a high-value crop that is cultivated on all continents, domesticated from a wild species in the genus *Zea*. It is an ancient allopolyploid, formed millions of years ago before the origin of the genus, by hybridization of two disparate parents. However, the identity of the parents has been unknown. We have discovered that all species of the genera *Zea* and *Tripsacum* are descended from this hybridization event, and the parents were probably African species, with one similar to modern *Vossia* (hippo grass) and *Urelytrum* (quinine grass), and the other possibly similar to modern *Rottboellia* or *Hemarthria*.

## Introduction

Cultivated maize (*Zea mays* ssp. *mays*) is an ancient allotetraploid, with two distinct subgenomes from unknown ancestors. The polyploid ancestry of the subspecies was initially suggested based on genetic evidence (1) and then confirmed with increasing amounts of molecular data (e.g., (2–4)) up to and including a full genome sequence (5, 6). The two subgenomes have differentiated and acquired distinct fates since being combined in a single nucleus exhibiting variation in expression patterns (7), fractionation (8, 9), neofunctionalization of homeologs (10) and many other aspects (e.g.(7, 11–15).)

The ancestral diploid genome donors are unknown, despite a wealth of genomic and phylogenetic data. The ancestral genomes appear to have last shared a common ancestor 13.6 ± 4.5 mya (16), with the polyloidization event (WGD) occurring after the divergence of *Zea* and *Sorghum* (16, 17). Using a set of low-copy nuclear genes, Estep et al. (16) hypothesized that the allopolyploidization event occurred before *Zea* diverged from its sister genus *Tripsacum*. However, that study included only *Z. mays* and *Tripsacum dactyloides* and did not include the other six species of *Zea* and 15 of *Tripsacum,*, all of which should bear the signature of the event.

The descendants of the *Zea-Tripsacum* genome donors are likely to be found, if they are still extant, among the group known informally as the awnless Andropogoneae (18). Andropogoneae (subfamily Panicoideae) is a clade of about 1200 species, which includes not only *Zea* and *Tripsacum*, but also sugarcane, *Miscanthus*, sorghum, and most of the species that dominate warm-season grasslands throughout the world (19–21). The flowers of most species are subtended by a bract (lemma) that ends in a long needle-shaped structure known as an awn, but in some taxa, such as maize and its relatives, the awn is absent. The other awnless Andropogoneae include about 185 species in 23 genera, most of which are native to Africa or Asia (22).

Because maize has a haploid chromosome number of *n*=10, authors in the pre-genomic era suggested that the ancestral genomes must have come from a species with *n*=5 (23), although only a handful of such species exist, most notably in the Asian genus *Coix*, which is also awnless. However, phylogenetic data do not support this hypothesis, and instead consistently point to species base number of 10, suggesting that the polyploidization event may have occurred between two *n*=10 ancestors (16, 24, 25), producing a plant with *n*=20; extensive genome rearrangement would have led to the current value of 10 (26). These studies have sampled only a few awnless species, raising the possibility that the closest relatives to *Zea* and *Tripsacum* have simply been missed.

To determine which species are descended from the allopolyploidization event that gave rise to *Zea* and *Tripsacum* and to identify species or clades whose ancestors might have donated one of the two genomes, we have used a combination of Sanger sequencing of nuclear genes, and Illumina sequencing of transcriptomic data. First, we added 35 species to the data produced by Estep et al. (16), which we refer to here as the 4-gene data set. Second, we combined data from the maize genome with transcriptomes for 20 species representing major Andropogoneae clades to generate gene trees placing each of the syntenic pairs of maize genes identified by Schnable and colleagues (14, 15). We then determined which of the awnless species were identified as sisters to one of the two paralogs. We take a doubly labeled gene tree as evidence for gene duplication. When the duplicated genes are dispersed throughout the genome, the pattern supports whole genome duplication. Finally, we constructed a phylogeny of whole chloroplast genomes (plastomes) to determine the maternal genome donor.

## Results

The 4-gene data show that the *Zea-Tripsacum* (hereafter, Z-T) allopolyploidization event occurred before diversification of the two genera (Figs. 1, S1). A clade composed of the awnless genera *Urelytrum digitatum* (quinine grass), *Vossia cuspidata* (hippo grass), *Oxyrachis gracillima* and *Rhytache rottboelloides* (hereafter the U-V clade), is well supported as sister to one set of Z-T paralogues, and excludes the other set, pointing to an ancestor of this clade as the donor of one Z-T (and hence maize) subgenome. The other set of paralogues may be sister to a clade including *Rottboellia* and *Hemarthria*, although support for this placement is weaker. *Coix* is unrelated to any of the other awnless taxa and no evidence places it close to *Zea* (Fig. 1, S1).

**Figure 1.**
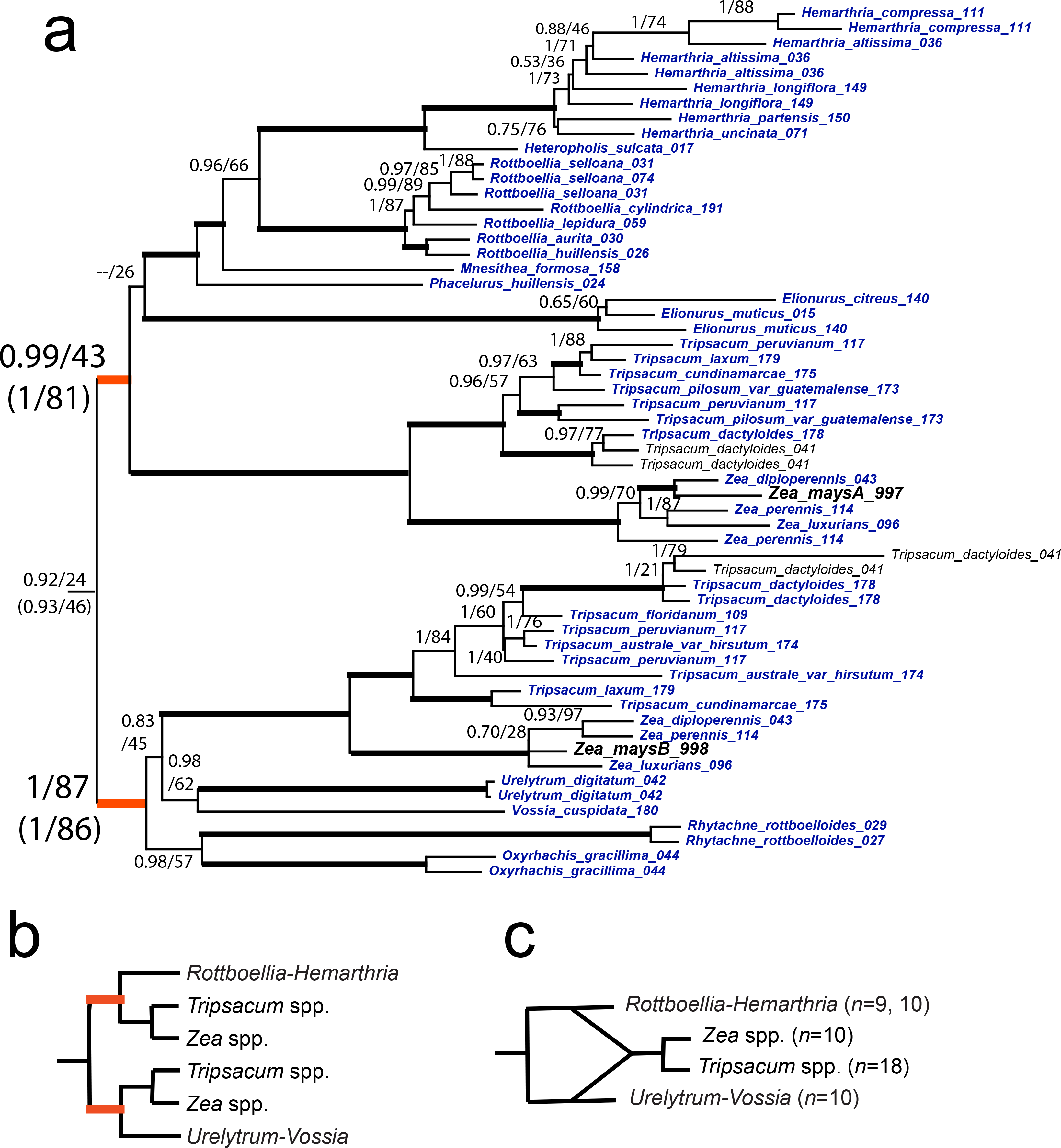
a. Phylogeny of *Zea*, *Tripsacum*, and the sisters to each of the subgenomes, based on four nuclear genes, concatenated; for full tree, see Figure S1. Numbers above branches are Bayesian posterior probability/maximum likelihood bootstrap. Heavy branches indicate support at both PP>0.95 and ML bootstrap >90. Branches highlighted in orange are those supporting the identity of the subgenomes. Support values in parentheses are those from trees with *Elionurus* removed. Blue lettering = new sequences, this study. b. cartoon summary of tree in a, excluding *Elionurus*. c. history of allopolyploidization inferred from tree in a.

The low support for the placement of the second Z-T clade led us to explore whether bootstrap values might reflect character conflict in the placement of *Elionurus*. Accordingly, we omitted the sequences from *E. tripsacoides* and *E. muticus*; in these analyses the bootstrap values placing Z-T clade 2 sister to the *Rottboellia-Hemarthria* clade were noticeably higher (82 vs. 42), suggesting strong signal connecting the latter two groups (Fig. 1).

To test the apparent sister relationship of the U-V clade to Z-T, we generated transcriptomic data and focused particularly on gene trees that included known syntenic pairs of paralogous genes (syntelogs) as defined by (14, 15). These are pairs of genes hypothesized to result from the allopolyploidization event. To mitigate the effect of missing data, we examined only gene trees that included, at a minimum, sequences from *Zea*, *Tripsacum*, *Urelytrum* and *Vossia*. (RNA was not available for *Oxyrachis* or *Rhytachne.)* In total, 2,615 gene trees met these conditions. Because some of these trees included more than one pair of paralogs, 3,351 maize paralog pairs were queried. Starting with a paralog (the target paralog) of each pair and working down the tree, clades were identified that either a) included a taxon other than *Zea* or *Tripsacum* (Fig. 2A), or b) include the sister paralog (Fig. 2B), and were supported by a bootstrap value (BSV) of at least 5 (orange branches, Figs. 2A and 2B). This procedure, implemented in PhyDS (see methods), identified 2,842 clades of paralogs. The low BSV was designed to include as many paralogous pairs as possible. Subsequent analyses then increased the BSV cutoff in increments of 5, up to 100. We then counted the numbers of clades that fit the pattern in Figs. 2A or 2B. If we found the 2B pattern, we counted clades in which X was *Urelytrum*, *Vossia*, or both, or another member of Andropogoneae. After filtering for taxa that were clearly misplaced based on the species phylogeny, we found that most commonly a *Zea mays* paralog was sister to its syntelog (1880 BSV 5 to 131 BSV 100, Table S3, Figs. 2A, C). The second most common relationship was a maize paralog sister to *Ureltyrum/Vossia* (595 BSV 5 to 7 BSV 100, Table S3, Figure 2B, C). Fewer clades found a maize paralog being sister to *Rottboellia/Hemarthria* (123 BSV 5 to 1 BSV 100, Table S3, Fig. 2C).

**Figure 2.**
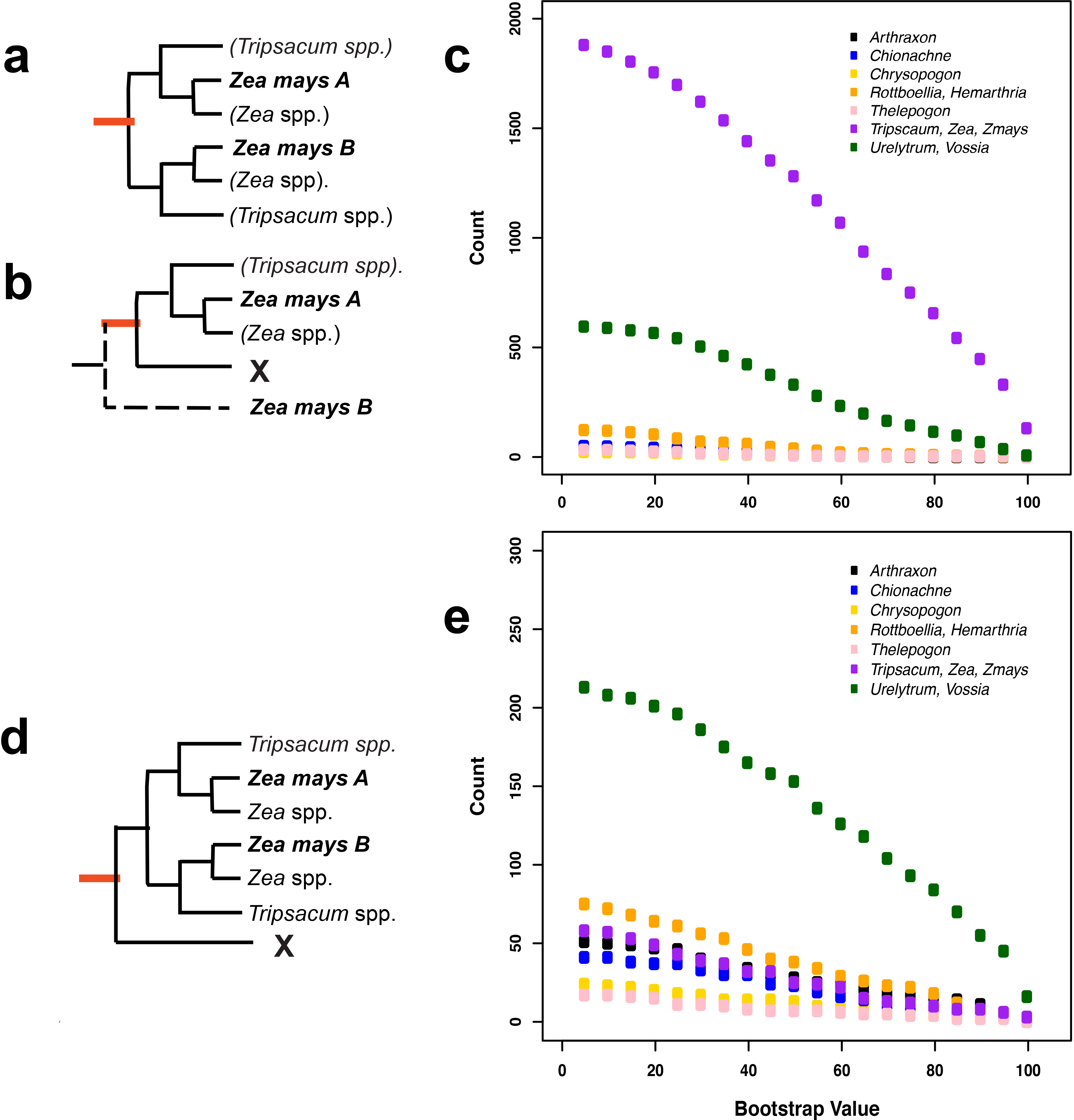
Phylogenomic identity of *Zea mays* paralogs. Gene trees identify the putative relationships (and ancestral identity) of maize paralogs. Queries were made to determine if Z-T paralogs were more closely related to each other than other taxa (A) or one paralog was more closely related to a different taxon than its sister paralog (B). Results filtered for bootstrap values (BSV) demonstrate that Z-T paralogs sister to each other (C, purple) is the most common, with U-V genes being sister to a Z-T paralog (C, green) the second most common pattern. Taxa identified as sister to a Z-T paralog clade (defined by D, seen as purple in C) were overwhelming from the U-V clade (E).

Trees were queried further for the 1,880 instances (940 clades, from the BSV 5 cutoff) in which maize paralogs were sister to each other, to determine the most common taxa sister to the pair of maize paralogs (pattern in Fig. 2D). The most common sister taxa were *Urelytrum, Vossia*, or both (213 BSV 5 to 16 BSV 100; Table S4; Figure 2E), and the second most commonly identified taxa were *Rottboellia/Hemarthria* (75 BSV 5 to 3 BSV 100; Table S4; Figure 2E).

Subgenome identities were mapped onto the maize genome (B73, annotation 6a) to visualize the distribution of parental subgenomes across the genomic landscape (Fig. 3). Concentrations of *Urelytrum/Vossia*-like genes were found in chromosomes 1, 2, 3, 5, 6, and 7. These distributions coincide with syntenic blocks, suggesting the conserved synteny of parental subgenomes even during the reorganization of the maize genome post polyploidy.

**Figure 3.**
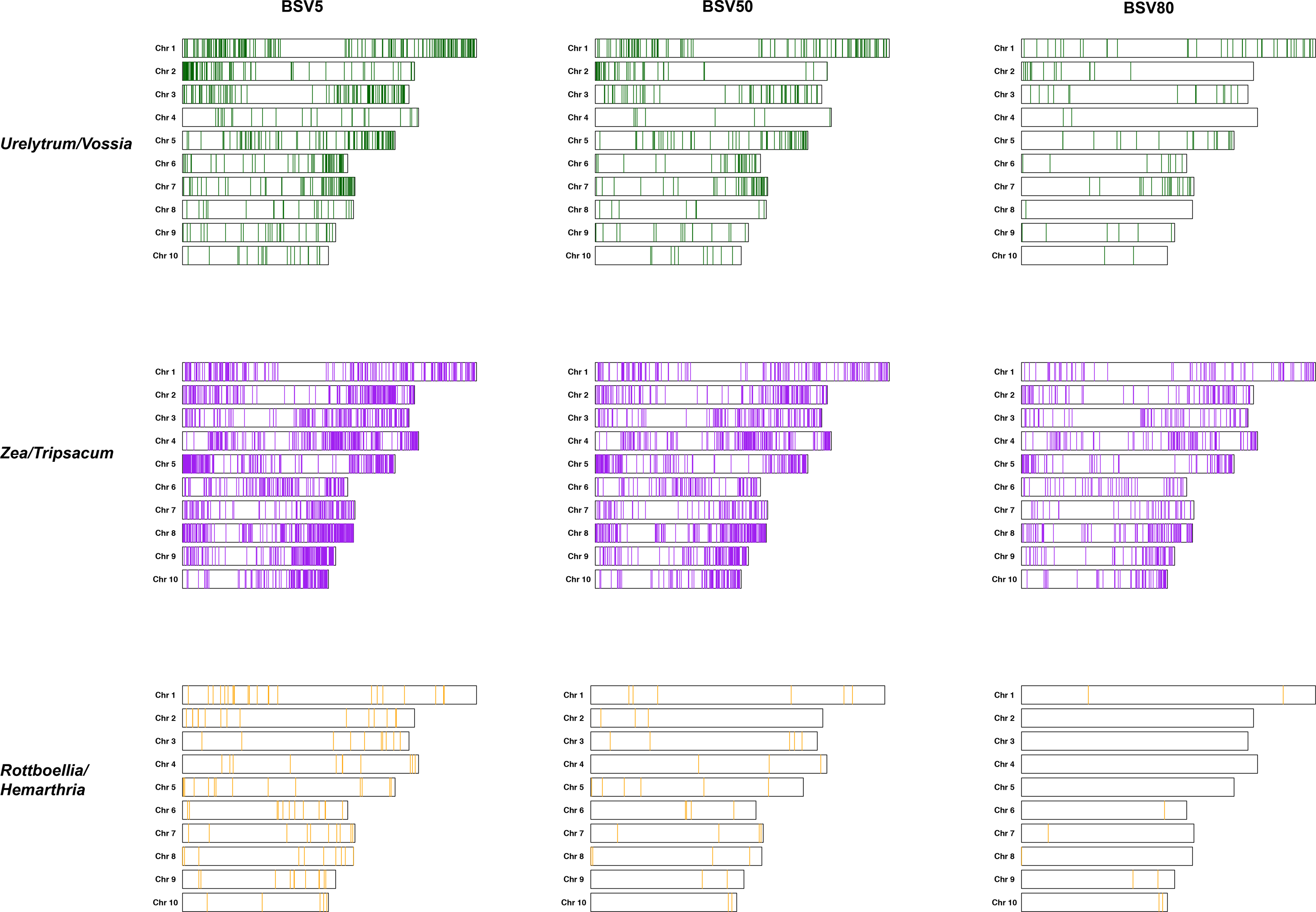
Subgenome identity of mapped genes. Genes were mapped to position across maize genome based on annotation information and assignment to subgenome identity based on transcriptome data and filtered by bootstrap values (BSV). Results suggest that portions of chromosomes 1,2,3,5, 6, and 7 are likely of U-V ancestral origin. Much of the genome demonstrates a close relationship between Z-T paralogs, though they are, in part, found in regions of low U-V identity. The *Rottboellia/Hemarthria-like* paralogs are few and do not have identifiable syntenic regions.

We also generated a chloroplast phylogeny of the Andropogoneae (Figs. 4, S2), which tracks the maternal genome. The Z-T clade was recovered with strong support, showing relationships similar to those in the nuclear trees, although we found some local disagreement between the plastome and the nuclear genes. However, the Z-T clade did not have any sister clades. While a monophyletic U-V clade was recovered, it and the Z-T clade were successive sisters to the remainder of the tribe. The *Rottboellia/Hemarthria* clade was only distantly related.

**Figure 4.**
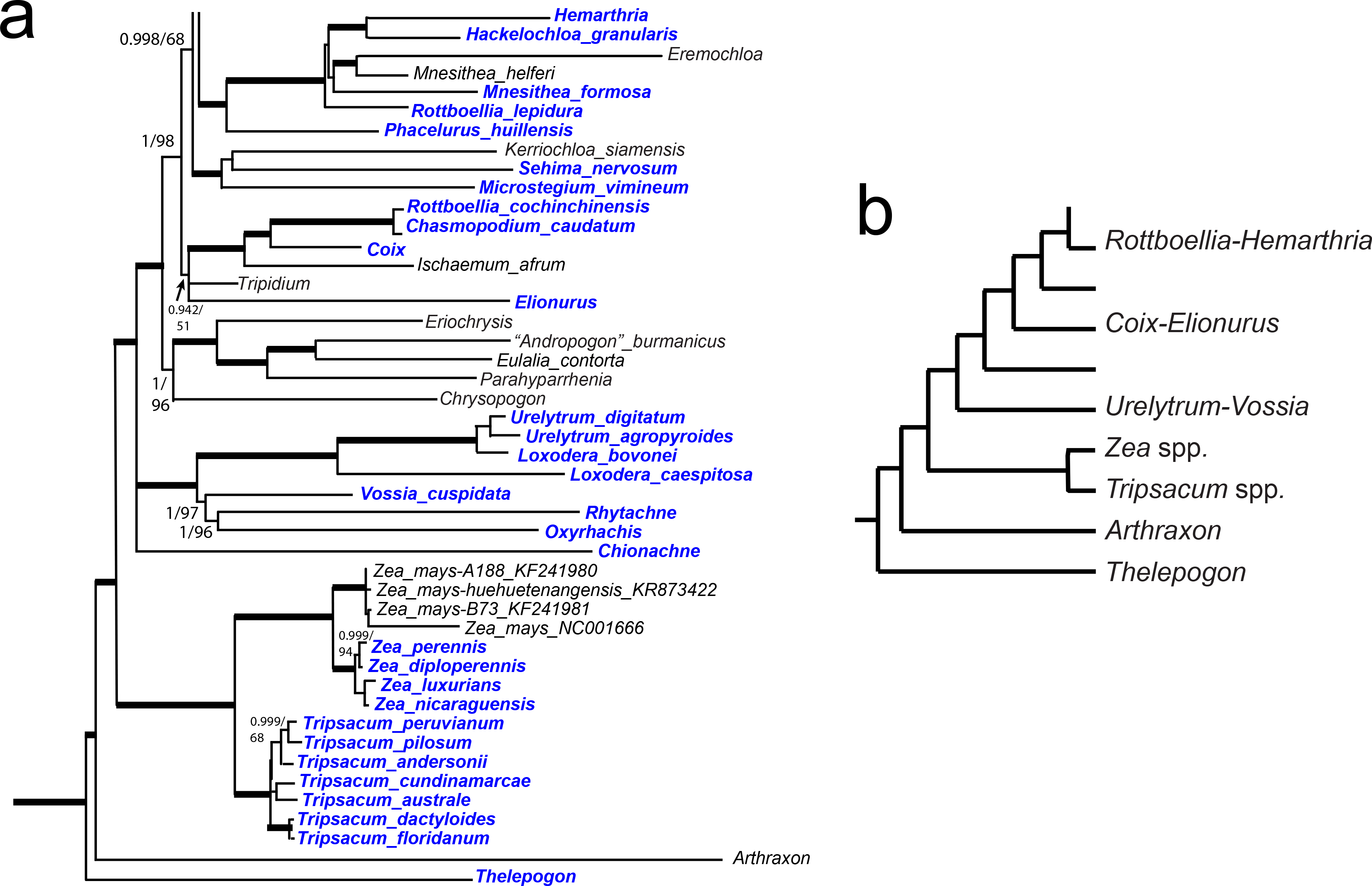
Phylogeny of *Zea*, *Tripsacum*, and their close relatives, based on complete plastomes; for full tree see Figure S2. No close relatives of the *Zea-Tripsacum* clade were identified, suggesting that either chloroplast lineage is unique to maize with no extant relatives or they have not been sampled.

## Discussion

Our data show that the ancestor of the U-V clade contributed extensive genomic material to the polyploidization event that characterizes the MRCA of *Zea* and *Tripsacum*, based on evidence from the 4-gene tree and from gene trees from the transcriptomes. An attractive hypothesis is that the U-V ancestor provided an entire genome, although evidence for this is obscured by subsequent rearrangements, gene evolution, and possible chromosomal conversion (27). The chloroplast phylogeny shows that the U-V clade is not the seed parent, but rather must have been the pollen donor.

The maize paralogs derived from U-V define some syntenic blocks and correlate with some, but not all, of the previously identified maize subgenomes (14, 15). In particular those on chromosome 1, 3, and 5 along with portions of 2, 6, and 7, suggest a close relationship to the UV-like parental genome (Fig. 4). However, there are exceptions. Instances of Z-T paralogs being sister to each other (Fig. 2 A, C (purple) and Fig. 3) are widespread across the genome, although they are concentrated in regions with fewer U-V-like paralogs (e.g. portions of chromosomes 2, 4, 8, 9, 10). Even where paralogous Z-T clades are sister to each other, the Z-T clade is often sister to U-V taxa (i.e., ((Z-T,Z-T),U-V)); Fig. 2 D, E). This is the expected pattern if these regions were derived from the U-V parental genome (i.e., ((Z-T,U-V),Z-T)), but the outgroup Z-T copy was converted by the UV-like copy (homeologous exchange with replacement; (27)). This possible mechanism is also consistent with the chloroplast tree (Fig. 4), in which the U-V clade is not sister to Z-T.

Members of the U-V clade have chromosome numbers in multiples of 10, whereas those in *Rottboellia/Hemarthria* have either 9 or 10 (Fig. 1C). *Tripsacum* has a base number of 18. The phylogenetic pattern suggests that the ancestral allopolyploidization event was the result of a cross between two parents with *x*=*n*=9 or 10, leading to a polyploid plant with *n*=19 or 20. We postulate a few chromosomal rearrangements soon after polyploidization leading to n=18, as seen today in *Tripsacum*, and then further rearrangements leading to *n*=10 in *Zea* (28). This hypothesis could be tested by comparing the *Tripsacum* genome to that of *Vossia* or *Urelytrum*, none of which are currently publicly available. We expect less rearrangement than in the *Vossia-Zea* comparison.

The data could be interpreted as supporting previous suggestions that maize is a segmental allotetraploid. In this model, some parts of the parental genomes recombine and undergo tetrasomic inheritance at the time of polyploidization, whereas others remain distinct and retain the evidence of their ancestry. This hypothesis was tested decades ago by Gaut and Doebley (4) using the handful of putative paralogues available at the time. We re-examined their data and found that, although most of their loci turn out not to be products of the allopolyploidy event, their conclusion is supported by the more extensive data now available.

Our data also show that the Z-T polypoidization event likely involved African ancestors. Few of the awnless species are native to the New World, and of those, none are closely related to *Zea* or *Tripsacum. Elionurus tripsacoides* is one such species, but it is more closely related to its African congener *E. muticus* than to any of the Z-T species or their relatives. While such a transatlantic distribution is surprising, it is not unheard of. Tetraploid cotton is well documented as a allopolyploid between an Old World and a New World species (29).

Identification of the paternal genome donor of *Zea* and *Tripsacum* may provide new insights into the genetics and physiology of the species. For example, several of the Z-T relatives are species of moist environments. *Vossia* grows in standing water, *Rhytachne* and *Oxyrhachis* in swamps, although *Urelytrum* occurs in open woodlands. This history could explain why maize is less drought tolerant than the related species sorghum (e.g., (30, 31)), but could also identify genomic regions that confer tolerance to waterlogged soils.

## Materials and Methods

### Taxon selection

Taxa were selected to encompass the diversity of *Tripsacum* and *Zea* and the majority of genera within Andropogoneae. Species names, voucher information, and GenBank accession numbers for the 4-gene data, transcriptome data, and plastome data are listed in Supplemental Tables 1-3, respectively. All data matrices and tree files are deposited at http://www.dryad.org, accession number xxx.

### DNA and RNA Isolation and Quantification

DNA was isolated using a modified CTAB protocol adapted from Doyle and Doyle (32) from live and silica dried leaves. DNA quantity was checked using a Nanodrop (ThermoFischer Scientific, Waltham, Massachusetts, USA) and a Qubit 2.0 (ThermoFischer Scientific, Waltham, Massachusetts, USA). Double-stranded DNA concentrations were used to determine starting material for library preparation. DNA quality controls required a 260/280 of 1.7 - 2.0 for isolation to be accepted.

RNA was isolated from ~0.5 g of young leaf tissue including unexposed leaf collected in midmorning from ideal greenhouse conditions. Samples were frozen in liquid nitrogen immediately upon collection. Samples were then crushed in an RNAse-free mortar with pestle in additional liquid nitrogen. Approximately 600 μL of TRIzol (Life Technologies) was added to each sample and ground with the sample. Samples were incubated at room temperature for 5-10 minutes. Chloroform was added in a 1:2 ratio (chloroform:TRIzol), and samples were vortexed and incubated at room temperature for 10 minutes. Samples were then spun at 12,000 x g for 15 minutes at 4C. The aqueous layer was transferred to a new tube and 1:1 of nuclease-free water was added. RNAse-free isopropanol was then added in a 1:1 and mixed by inversion. Samples were incubated at room temperature for 10 minutes and then spun at 12,000 × g for 15 minutes at 4C to pellet out the nucleic acids. The pellets were washed with fresh, RNAse-free 80% ethanol and spun at 12,000 × g for 5 minutes at 4C. The 80% ethanol was removed and the pellets were air dried for 10 minutes. Samples were suspended in 30-50 μL of nuclease-free water. Samples were DNAsed using DNAse I from the DNA-Free DNA Removal Kit (AM1906, ThermoFischer Scientific, Waltham, MA, USA) per manufacturer’s protocol. Samples were split into aliquots to minimize freezing and thawing of any one sample.

RNA quality was measured on an Agilent 2100 Bioanalyzer using an Agilent RNA 6000 Pico chip (Agilent Technologies, Inc., Santa Clara, CA, USA). Samples were diluted 1:100 prior to running the analysis. RIN scores were used to determine quality of RNA with a soft cutoff of 8.0 and a hard cutoff of 7.0.

### Four-locus Data Sequencing and Processing

Data for the four-locus phylogeny was obtained in the same manner as (16) with some minor alterations. Trees were estimated using RAxML v.8.0.22 (33) under the GTR + gamma model with 500 bootstrap replicates. A Bayesian inference (BI) phylogeny was reconstructed using MrBayes v.3.2.6 (34) with two independent runs with four chains—one cold and three heated—with six substitution types under the GTR+gamma evolutionary model sampling every 1,000 for 5,000,000 generations. Convergence was verified using potential scale reduction factor (PSRF) values calculated by MrBayes.

### Illumina Library Preparation and Sequencing

DNA samples were sheared using a Covaris S220 (Covaris, Inc., Woburn, MA, USA) with a target size of 300-500 bp at either the University of Missouri DNA Core Facility or the Donald Danforth Plant Science Center. Sequencing libraries were made using the NEBNext Ultra DNA Library Prep Kit for Illumina (New England BioLabs, MA, USA) or the Nextera DNA Library Prep Kit for Illumina (Illumina, Inc., San Diego, CA, USA) per manufacturer’s instruction. Libraries were size selected for 300-400 base pair insert sizes. Sequencing was conducted at the University of Missouri DNA Core Facility, the New York University School of Medicine Genome Technology Center, the University of Georgia Genomics Facility, or the University of Illinois Roy J. Carver Biotechnology Center. Single-end 100 base pair runs on a HiSeq2500 (UM), a paired-end 150 base pair rapid run on a HiSeq2500 (NYU), a paired-end 150 base pair run on a NextSeq500 (UGA), and a paired-end 250 base-pair rapid run on a HiSeq2500 (UI) were used to sequence libraries.

Total RNA was enriched using the Magnetic mRNA Isolation Kit (S1550, New England BioLabs, Ipswich, Massachusetts). In the heated elution step, samples were heated at 94 ° for 810 minutes to target 400-500 base pair fragments. mRNA libraries were made using the NEBNext Ultra Directional RNA Library Prep Kit for Illumina (New England BioLabs, Ipswich, Massachusetts, USA) per manufacturer’s protocol with Agencourt AMPure XP bead (Beckman Coulter, Indianapolis, Indiana, USA) size selection targeted for 400-500 bp insert sizes. Libraries were quantified using an Agilent 2100 Bioanalyzer with a High Sensitivity DNA Analysis chip (5067-4626, Agilent, Santa Clara, California, USA). Libraries were diluted to a final concentration of 10 nM and pooled in equal volumes in sets of 5 and to be sequenced using 150 bp paired-end sequencing at the University of Wisconsin Biotechnology Center on an Illumina HiSeq2500 or a set of 9 (including samples not presented in this manuscript) to be sequenced using 150 bp paired-end sequencing on an Illumina NextSeq500 at the Georgia Genomics and Bioinformatics Core.

### Chloroplast Genome Assembly, Annotation, and Tree Estimation

Chloroplast genomes were assembled using the Fast-Plast method (35). Raw reads were trimmed using Trimmomatic v.0.36 (36) with adapter trimming of appropriate adapters (Nextera or NEBNext), a seed mismatch of 1, a palindrome clip threshold of 30, and a simple clip threshold of 10. Reads were further filtered using a sliding window of 10 basepairs with a minimum average quality score of 20 and a minimum length of 40. Reads were then mapped to a set of Andropogoneae chloroplast genomes identified from GenBank, which was updated as plastomes were finished, using bowtie2 v.2.1.0 (37) under the “very-sensitive-local” parameter set. Mapped reads were assembled using SPAdes v.3.9.0 (38) with the “only-assembler” option and k-mer sizes of 55, 69, and 87 for sequencing run with reads of 100 bp and 55, 87, and 121 for runs with reads larger than 100 bp. Assemblies were filtered based on coverage estimated by SPAdes to remove extremes of coverage in contigs less than 1000 bp. The program afin (in FastPlast (35)) was used to attempt contig fusion and extension using the full, trimmed read set with three iterations. The first iteration consisted of 150 search loops, an initial contig trim of 100 bp, a minimum overlap 20 bp, a minimum coverage of 2, and a max extension length of 0.75 × the max read length for the sample. The second and third iterations consisted of 50 search loops, an initial contig trim of 100 bp, a minimum coverage of 1, and differed with a minimum overlap of 15 and 10 bp, respectively. If the result was a single contig, the assembly sequence_based_ir_id.pl and orientate_plastome_v.2.0.pl scripts were used to identify boundaries of the small single copy, large single copy, and inverted repeat regions of the chloroplast genomes (35). If an afin run did not result in a full plastome, contigs were assembled manually in Sequencher v.5.3 (Gene Codes Corporation) using *Schizachyrium scoparium* (NC_035032.1) to scaffold as described in McKain et al. (39). Gaps were closed through *in silico* plastome walking if possible, but otherwise filled using N’s. A coverage analysis as implemented in Fast-Plast was done for each plastome to identify regions of low quality assembly. These were investigated manually and adjusted based on read coverage when necessary.

Chloroplast genomes were annotated using Verdant (40). Annotations were verified using GenBank validation (https://github.com/mrmckain/Verdant_Utilities/). If any genes were found to be missing from the annotation or seemingly pseudogenized, the assemblies were verified by mapping reads as above and visual inspection in Sequencher. If an alternative assembly was found to be more supported and provided sequence for a functional gene, the assembly was changed and the plastome reannotated. Final annotations are available in Verdant (verdant.iplantcollaborative.org) and GenBank (XXX-XXX).

Chloroplast genomes were split into their subcomponent regions: LSC (large single copy), SSC (small single copy), and IR (inverted repeat) using a Perl script (https://github.com/mrmckain/Genome_Skimming_Utilities). These regions were aligned individually using MAFFT v.7.245 (41) using high speed default parameters. Once aligned, these regions were concatenated using a Perl script (https://github.com/mrmckain/Genome_Skimming_Utilities). The best base pair substitution model was determined to be GTR + gamma + I using the Aikake Information Criterion (AIC) in jModelTest2 v.2.1.5 (42). The total data set had 173 taxa (S1) and included taxa from GenBank, Arthan et al. (43), Welker et al. (44), and newly sequenced plastomes from this study.

A maximum likelihood (ML) phylogeny was reconstructed using RAxML v.8.0.22 (33) under the GTR+GAMMA+I evolutionary model with 500 bootstrap replicates. Bayesian inference (BI) phylogenies were reconstructed using MrBayes v.3.2.6 (34) with six independent MCMCMC runs with four chains—one cold and three heated—with six substitution types under the GTR+GAMMA+I evolutionary model sampling every 1,000 for 1,000,000 generations. Convergence was verified using PSRF values calculated by MrBayes.

### Transcriptome Assembly, Annotation, and Gene Tree Estimation

Raw transcriptome reads were assembled using Trinity v.r20140413 (45). In-silico normalization, as implemented in Trinity, was used to reduced read coverage over 50×, followed by read cleaning using Trimmomatic v.0.30 (36) using a sliding window of 10 bp with a minimum average quality score of 20. When libraries were made using the NEBNext Directional Kit, the stranded option was used in Trinity. Other parameters were defaults. After assembly, the “align_and_estimate_abundance.pl” script packaged with Trinity was used to estimate read abundance and the FPKM for each script using RSEM v.1.2.1 (46). Transcripts were filtered using the “filter_fasta_by_rsem_values.pl” packaged with Trinity and set to a minimum isoform percentage value of 1.00.

Once transcripts were filtered, coding sequence was estimating using Ref-Trans (39) by BLASTing the transcripts against a database of CDNA from all available PACMAD genomes in Phytozome 10 (https://phytozome.jgi.doe.gov/pz/portal.html) using tblastx with an e-value cutoff of 1e-10. A reciprocal 85% overlap of a transcript and a database sequence was required to proceed to annotation. Best matches were used to guide translations with GeneWise (part of the Wise v.2.2.0 package; (47) under default parameters. For each translation, the longest identified reading frame was used in subsequent steps. Annotated sequences were identified to orthogroups using OrthoFinder v.0.2.0 (48) with default parameters. Orthogroups were filtered so they had at least five representative taxa. Amino acids for each orthogroup were aligned using PASTA v.1.6.3 (49) with the “auto” parameter setting. PAL2NAL v.14 (50) was used to produce a codon alignment from the amino acid alignment under default parameters. Orthogroups were further filtered by sorting aligned orthogroups into clusters with a Hamming distance threshold of 0.25 using a Perl script (core_split_alignments_distance_v1.5.pl). These aligned clusters were used to reconstruct gene trees. Gene trees were estimated using RAxML v.8.0.22 under the GTR + gamma model with 500 bootstrap replicates. Trees were rooted to either a *Dichanthelium oligosanthes* (v.1.0; (51)), *Panicum virgatum* (v1.1, DOE-JGI, http://phytozome.jgi.doe.gov), or *Setaria italica* (v.2.1; (52)). The best trees (highest likelihood) and their bootstrap values were kept for downstream analyses.

### Identification of Maize Subgenomes

Paralogs were identified through synteny analysis from the *Zea mays* genome (annotation Zmays_284_6a) following the methods of Schnable and colleagues (7, 14, 15) as implemented in CoGe (53–56). These putative paralogs were used to query gene trees estimated from transcriptome and genome data. The trees with bootstrap values were filtered so that only those with a pair of maize paralogs and at least one transcript from either *Urelytrum* or *Vossia* was present.

The package PhyDS (https://github.com/mrmckain/PhyDS) was used to identify the putative evolutionary origin of identified maize paralogs. For each paralog, the PhyDS algorithm starts at the representative tip of the tree and moves to the ancestor node. In each iteration, descendants of the current node are identified. PhyDS was parameterized so that this process was iterated until either 1) a taxon was encountered that was not from either *Zea* or *Tripsacum* or 2) the sister paralog (syntelog) to the current target paralog was found. When either of these conditions was met, the bootstrap value was recorded for the current node as well as the descendent genes identities. If the *Zea mays* sister paralog was encountered, then the sister lineage was designated as *Zea mays* to differentiate these instances. Results were filtered using scripts included with PhyDS to remove instances where outgroups *(Dichanthelium, Panicum*, or *Setaria), Arundinella, Sorghum, Ischaemum, Miscanthus, Saccharum, Themeda, Bothriochloa, Dichanthium, Andropogon*, or *Schizachyrium* were present in the descendants of the ancestral node. These were chosen to be excluded based on known relationships in Andropogoneae (43, 44, 57) and the chloroplast genome phylogeny presented in this study. Andropogoneae nuclear gene trees are known for extremely short branches potentially due to a rapid diversification (16) which can impede identification of relationships among paralogs and orthologs but can be lessened with the removal of suspect topologies. Filtered results were binned based on minimum BSV thresholds starting at BSV 5 and increasing by increments of five until BSV 100. Results were further filtered to identify clade composition. If only *Zea* or *Tripsacum* were identified, this was counted as a *Zea/Tripsacum*. If *Zea* and *Tripsacum* were identified along with one of the following: 1) only *Arthraxon*, 2) only *Chionachne*, 3) only *Chrysopogon*, 4) *Rottboellia* and/or *Hemarthria*, 5) only *Thelopogon*, or 6) *Urelytrum* and/or *Vossia*, the respective taxa were counted as being more closely related to the target paralog.

For cases where the *Zea mays* paralog was found to be more closely related to its sister paralog than other taxa, we further investigated what lineage was sister to this clade. We used paralog pairs identified from the BSV 5 binned results to query gene trees with PUG (39) under default parameters and the chloroplast tree used as the species tree. We used both the “Gene_Tree_Results” and the “Gene_Trees_Pairs_Bad_Results” files from PUG for further analysis. These files have the same format, except the “Gene_Trees_Pairs_Bad_Results” file represents genes trees that do follow the topology of the given species tree. Since we used PUG for its ability to give us the sister lineage taxon composition to a clade consisting of paralogs and their MRCA, the comparison to the species tree did not matter. We then filtered these results in the same manner as above based on taxon composition and BSV.

## Acknowledgements

This project was supported by NSF grants DEB-11456884 and DEB-1457748 to EAK. Pre-publication data for the *Panicum virgatum* genome was provided by the Department of Energy Joint Genome Institute. Collecting in Tanzania by STM was done in collaboration with the Missouri Botanical Garden through a permit from COSTECH. We thank James Schnable for pre-publication access to an updated list of paired syntenic loci in the maize genome. We also thank Cassiano A. D. Welker for DNA of *Schizachyrium sanguineum* for the plastome sequence, and Trevor Hodkinson for DNA of *Polytoca* and *Hemarthria* for the 4-gene tree. Shao Ying in the lab of Thomas Brutnell extracted several of the RNA samples used for transcriptomes.

## Author contributions

Conceptualization, Methodology: MRM, MCE, EAK. Software: MRM. Formal Analysis: MRM. Investigation: MCE, DJL, DMVD, JGH, JZ. Resources: RP, DJL, STM, DMVD, JZ, GC, BP, EAK. Data Curation: MRM. Writing - Original Draft: MRM, MCE, EAK. Writing - Review & Editing: All authors. Visualization: EAK. Supervision: MRM, MCE, EAK. Project Administration and Funding Acquisition: EAK.

## Supplemental Information

Table S1. Voucher and Genbank numbers for newly added taxa in 4-gene tree. All others from Estep et al. (2014).

Table S2. Transcript assembly information for transcriptome data.

Table S3. Voucher and GenBank accession numbers of plastome data.

Table S4. Numbers of clades with the given taxon sister to a Zea-Tripsacum paralog. BSV = ML bootstrap support value for the relevant clade.

Table S5. Numbers of clades with the given taxon sister to the clade including both Zea-Tripsacum paralogs. BSV = ML bootstrap support value for the relevant clade.

Figure S1. Phylogeny of Andropogoneae based on four nuclear genes, concatenated. Numbers after taxa are accession numbers. Sequences from the same accession number are different cloned sequences from the same plant. Numbers above branches are Bayesian posterior probability/maximum likelihood bootstrap. Shaded boxes delimit subtribes, following Kellogg (2015).

Figure S2. Phylogeny of Andropogoneae based on complete plastomes. Numbers above branches are Bayesian posterior probability/maximum likelihood bootstrap.

